# Bwa, an ortholog of alkaline ceramidase-ACER2, promotes intestinal stem cell proliferation through pro-inflammatory cytokine signaling in *Drosophila melanogaster*

**DOI:** 10.1101/2024.11.26.624044

**Authors:** M. Mahidur Rahman, Chloe Kraft, Collin Clark, Rebekah J. Nicholson, Marco Marchetti, Emily Williams, Chenge Zhang, Will L. Holland, Scott A. Summers, Bruce A. Edgar

## Abstract

Sphingolipids, including ceramides, are an important component of high-fat diets. These molecules can regulate fatty acid oxidation and intestinal stem cell proliferation, predisposing the gut to tumorigenesis. However, the molecular mechanisms involved in ceramide metabolism-mediated intestinal stem cell (ISC) proliferation and tumorigenesis are poorly understood. To understand how changes in sphingolipid metabolite flux affect intestinal stem cells, we manipulated the activities of each of the enzymes of the ceramide synthetic pathway using cell type-specific over-expression or depletion of the corresponding mRNAs in each intestinal cell type of the *Drosophila* midgut. We documented cell-autonomous and non-cell-autonomous effects, including alterations in cell size, number, differentiation, and proliferation. In our screen, the altered expression of several ceramide metabolism enzymes led to changes in ISC proliferation, cell sizes, and overall cellularity. Among other genes, over-expression of ceramidase homolog, *Brain washing (bwa)* in gut enteroblasts (EB) increased EB cell size and caused a non-cell-autonomous, 7-8-fold increase in ISC proliferation. Our analysis confirmed previous reports that *bwa* does not have ceramidase activity, and lipidomic studies indicated that *bwa* increases the saturation status of sphingolipids, free fatty acids, and other lipids. The pro-proliferative effects of *bwa* could be counter-acted by depleting a serine palmitoyltransferase, *Lace*, or a sphingosine acyltransferase, *Schlank*, which are needed for ceramide synthesis, or by co-expressing a ceramide desaturase enzyme, *ifc*, indicating that increased saturated ceramides were causal for ISC proliferation and the disruption of gut homeostasis. Accumulating saturated sphingolipids and fatty acids induced inflammatory signaling in the gut, and activated ISC proliferation through the pro-inflammatory cytokines, Upd3 and Upd2. We propose that saturated sphingolipids promote ISC proliferation through pro-inflammatory pathways.

## INTRODUCTION & BACKGROUND

A high-fat diet is a known risk factor for many adverse health outcomes, including type II diabetes, inflammatory bowel disease, and colorectal cancer. Although it is still being determined how the different metabolites in lipid-rich diets affect cellular functions, increased sphingolipid signaling is one major effect of high-fat diets^1^ that is believed to be responsible. One of the common components of sphingolipid – ceramides^2^ can regulate cellular processes such as fatty acid transport and metabolism, apoptosis, cell growth, and proliferation in addition to its structural functions in the lipid bilayers of the cellular organelles^3,4^. Ceramides/ sphingolipid metabolites have been associated with cellular signaling mechanisms in both normal and disease cell physiological states^5,6,7,8^.

Ceramide, the abundant precursor to most sphingolipids, typically comprises an 18-carbon sphingosine base linked to a long-chain fatty acid component^9^. Ceramides are synthesized by four major pathways: endogenous de novo biosynthesis, hydrolysis of more complex sphingolipids such as sphingomyelin into their constituents, the salvage pathway, and fatty acid oxidation^10^. Recent research has highlighted the impact of dietary sphingolipids on intestinal microbiota and gastrointestinal (GI) immune homeostasis. Sphingolipids provide structural integrity to the gut epithelial cells, regulate nutrient absorption, and act as receptors for microbial antigens and toxins. Sphingolipids such as sphingomyelin (SM), sphingosine (Sph), ceramide (Cer), sphingosine-1-phosphate (S1P), and ceramide-1-phosphate (C1P) are not only structural components of cell membranes but also modulate cellular processes like vesicle trafficking, proliferation, and apoptosis. These processes are essential for maintaining GI barrier function and immune response. One of the first enzymes in the ceramide biosynthetic pathway-*serine palmitoyltransferase long chain base subunit 1 and 2* (SPTLC1&2), which diverts dietary fatty acids and amino acids into the biosynthetic pathway, is an important regulator of intestinal stem cell homeostasis^11^. An imbalance in dietary sphingolipids has been linked to chronic inflammation and an increased prevalence of GI disorders^12^. Sphingolipid metabolism dysregulation correlates with the pathogenesis of metabolic diseases, including those affecting the GI tract. Bioactive sphingolipids like ceramide and S1P regulate cellular growth, differentiation, and apoptosis, which are critical for the physiological functioning of the GI system^13^. Consequently, sphingolipids and their metabolic enzymes are being explored as therapeutic targets and potential biomarkers for disease status. Their roles in inflammatory bowel disease (IBD) and colorectal cancer (CRC) is of particular interest, as sphingolipids regulate inflammation, cell signaling, differentiation, apoptosis, and survival, all of which are implicated in IBD and CRC^13^. A network of complex interactions exists between dietary sphingolipids, gut microbiota, and the host immune system. Ceramides, thus, have emerged as a central lipid species at the cross-section of metabolomics, nutrition science, cardiovascular health, aging, and cancer biology^14,15,16,17,18^. Understanding the mechanisms involved in this ceramide centric network of cellular interactions is vital for developing effective interventions for better gut health and therapeutic intervention.

While disease associations of enzymes responsible for ceramide/sphingolipid metabolism have been identified, discovering the mechanistic details and effects of those associations with cellular processes is an ongoing quest. Often, these end-point studies are hard to interpret - raising questions such as whether an alteration in a sphingolipid metabolite is an effect or a cause of the presented phenotype. Lack of tools to probe the impact of altered sphingolipid metabolism and easy readouts to infer the role of these metabolites in biological processes have complicated the subject even more. The *Drosophila* midgut presents an excellent system in which it is possible to probe gene functions in the different intestinal cell types, namely ISCs and their differentiated progeny: transient enteroblasts (EB), absorptive enterocytes (EC), and secretory entero-endocrine (EE) cells. With the availability of Gal4-based temporally controlled cell type-specific genetic drivers, we found that it was possible to dial up or down the expression of each gene encoding the enzymes in the ceramide synthetic pathways. After doing this, we scored various parameters of gut cell physiology, such as proliferation, differentiation, cell size, cellularity, and cell-autonomous and non-cell-autonomous effects. In this screen, an enigmatic gene called *brainwashing (bwa),* an enzymatically non-functional ortholog of the mammalian alkaline ceramidase (ACER2), emerged as one of the most potent drivers of ISC proliferation. Through lipidomic experiments and functional tests, we analyzed the nature of changes in sphingolipid metabolites after manipulating the Bwa function. Here, we present our observations on the impact of ceramide metabolism on *Drosophila* ISCs and their lineage.

## RESULTS

### Ceramide metabolism genes affect ISC functions and gut homeostasis

The enzymes involved in the synthesizing and degradation of sphingolipids and ceramides have been identified and characterized in some detail^19^. However, the functions of these genes in the physiology of different organs and cell types *in vivo* are less well known. To address the functions of the various ceramide metabolism enzymes in the gut, we used transgenic, temperature-sensitive, and cell-type-specific Gal4 drivers to express RNAi or overexpression (Open Reading Frame; ORF) constructs for each gene in the sphingolipid metabolic pathway. For cell type-specific expression of the ORFs/RNAis, we used the *esgG4 tubG80^ts^ Su(H)-GBE-Gal80* driver (for ISCs, designated as ISC-G4^ts^, *Su(H)-GBE-Gal4 tubGal80^ts^* (for EBs, designated as EB-G4^ts^), *Myo1A-Gal4 tubGal80^ts^* (for ECs, designated as EC-G4^ts^) , and *ProsV1Gal4 tubGal80^ts^* (for EEs, designated as EE-G4^ts)20,21^. In each case, the UAS-GFP transgene was co-expressed to mark the affected cells. As shown in Supplementary Figure 1 and Supplementary Figure 2 [BE1] [MR2] [MR3] [MR4] and summarized in Tables 1 and 2, genetic perturbation of the enzymes involved in ceramide metabolism in different cell types showed many strong, robust phenotypic alterations in the size, shape, and number of the cell types, and these changes were both cell-autonomous and non-cell-autonomous. For example, the overexpression of *lace^ORF^*to generate a gain of function of serine C-palmitoyltransferase, one of the first enzymes in the ceramide biosynthetic pathway (Figure 1 A), caused the cell-autonomous effect of increasing the number and size of the ISCs without affecting the other cell types. However, the overexpression of *lace^ORF^* in the EBs, ECs, and EEs had the obvious albeit relatively mild non-cell-autonomous effect of increasing ISC proliferation (Supplementary Figure 1, Supplementary Figure 2). Conversely, RNAi-mediated knock-down of lace in ISCs reduced ISC numbers, whereas *lace^RNAi^* expressed in ECs or EEs mildly increased ISC proliferation. Similarly, the expression of *schlank^ORF^*, which encodes a sphingosine N-acyltransferase, increased the number and size of ISCs and EBs with some loss of ECs and no effect on the EEs. Conversely, *schlank^RNAi^* expression reduced the number and size of the ISCs and EBs without affecting the ECs and EEs (Supplementary Figure 1, Supplementary Figure 2, and Tables 1 and 2). Other notable changes in the morphology, number, and proliferation pattern of the ISCs were affected by the alterations of the levels of the enzymes such as bwa, ifc, cdase, sk2, cpt1, snmase, ksr, and fatp1-all involved in the ceramide metabolic pathway, are summarized in Table 1 (Supplementary Figure 1, Supplementary Figure 2). Of these, *lace^ORF^, schlank^ORF^, ifc^RNAi^, glc-t^RNAi^, sk2^RNAi^*and *cpt1^RNAi^* increased ISC proliferation autonomously and non-autonomously. On the other hand, *ifc^ORF^, fatp1^RNAi^* and *fabp1^RNAi^* could induce ISC proliferation cell non-autonomously only when expressed in the enterocytes. It is unclear if some of these effects result from cell death induced by altering the levels of the ceramide metabolic enzymes. In cases of *bwa^ORF^, schlank^ORF^, ifc^RNAi^, snmase^RNAi^, cpt1^RNAi^*, and *fabp1^RNAi^* expressed in enteroblasts, however, there was no obvious cell death. Rather, the EBs were enlarged, suggesting that other cell signaling pathway(s) could induce the increased ISC proliferation cell non-autonomously (Supplementary Figure 1, Supplementary Figure 2, Tables 1 and 2). Given that only about 7-10 PH3+ cells are found in the midgut in the basal condition, it is difficult to identify a perturbation with confidence where the ISC proliferation is reduced. However, the knockdown of fabp1 in the ISCs showed a robust ten-fold decrease in ISC proliferation, suggesting a cell-autonomous requirement for fabp1 in the ISCs (Table 2, Supplementary Figure 2). Knockdown of the same fabp1 gene in the terminally differentiated ECs, and EEs, however, is associated with non-cell-autonomous ISC proliferation (Table 2, Supplementary Figure 2), suggesting an important but divergent role for this fatty acid binding protein in the ISC lineage.

**Figure 1.**
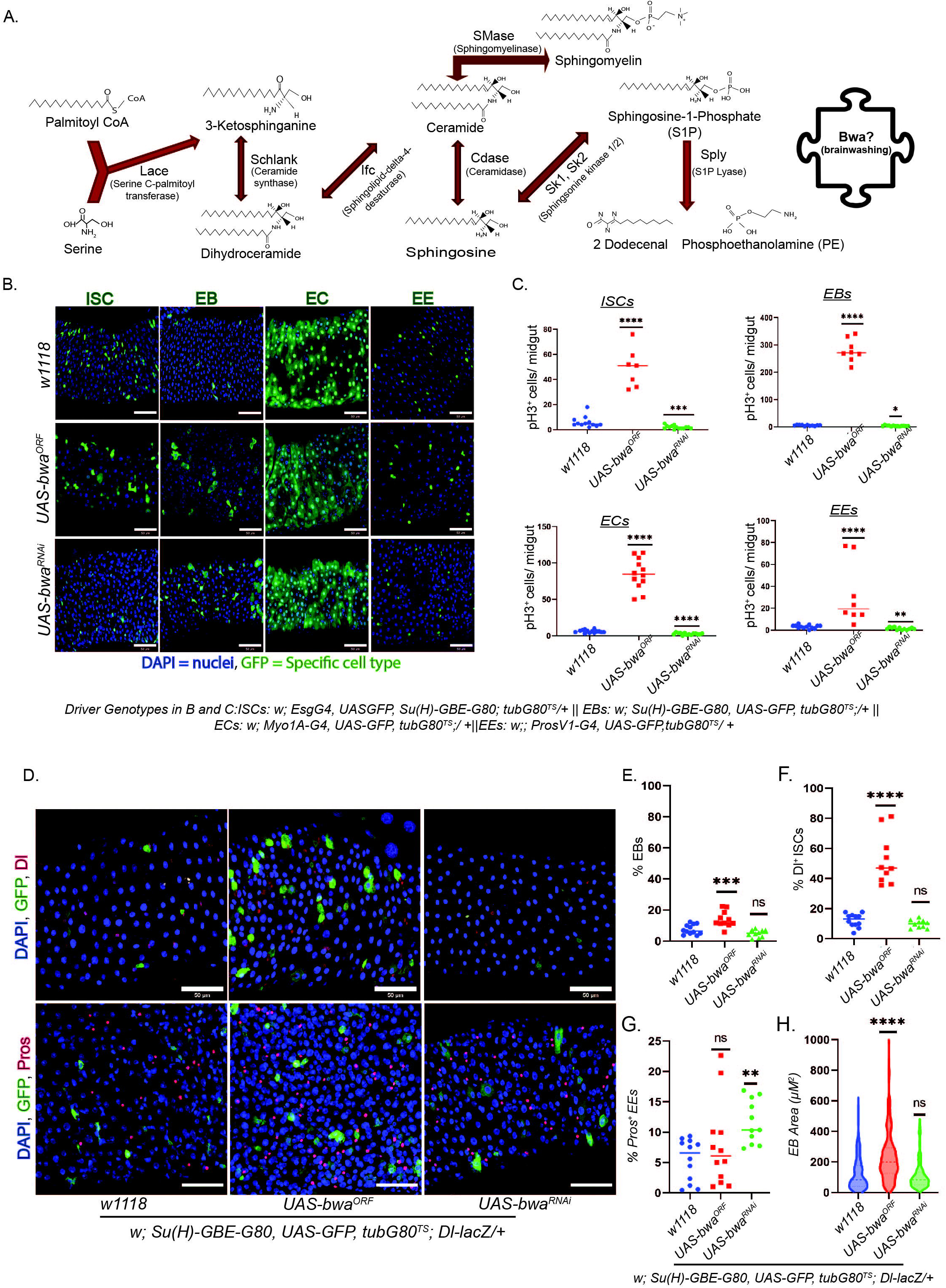
Brainwashing, Bwa, is an enigmatic enzyme that promotes intestinal stem cell proliferation and enteroblast growth. A. Cartoon showing the general ceramide metabolic pathway, including the chemical structures of the different sphingolipid species and intermediates. B. Images showing the different cell types in the posterior midgut with Bwa gain-of-function and reduced levels compared to controls at permissive temperatures for five days. C. Graphs cell-autonomous and cell-non-autonomous effects of Bwa levels on ISC proliferation. Images showing stem cell marker Dl+ (red), and enteroendocrine cell maker Pros+ (red) in the EBs (green) in EB-specific Bwa Gain of function and Loss-of-function conditions compared to control (w1118) conditions. E-G, graphs show the relative proportions of the EBs, ISCs, and EEs, while graph H shows the relative size (area) of EBs in in EB-specific Bwa Gain of function and Loss-of-function conditions compared to control (w1118) conditions. Scale bars represent 50 µM, * represents values e.g. *\* p< 0.05 and, ** p< 0.005, *** p< 0.0005, **** p< 0.00005*.

**Table 1:** Cell-autonomous morphological effects of over-expression or knockdown of enzymes involved in ceramide metabolism in the ISC lineage.

| **Perturbation** | **ISCs** | **EBs** | **ECs** | **EEs** |
| --- | --- | --- | --- | --- |
| *UAS-lace*<sup>ORF</sup> | Increased number and size | Unaffected | Unaffected | Unaffected |
| *UAS-lace*<sup>RNAi</sup> | Loss of ISCs | Loss of EBs | Unaffected | Misshapen, larger in size |
| *UAS-schlank*<sup>ORF</sup> | Increased number and size | Larger in size, no change in numbers | Some loss of ECs | Unaffected |
| *UAS-schlank*<sup>RNAi</sup> | Rounded ISCs | Unaffected | Unaffected | Larger in size |
| *UAS-ifc*<sup>ORF</sup> | Rounded ISCs | Unaffected | Unaffected | Unaffected |
| *UAS-ifc*<sup>RNAi</sup> | Larger in size | Larger in size | Unaffected | Reduced number |
| *UAS-cdase*<sup>RNAi</sup> | Reduced number | Reduced number | Unaffected | Unaffected |
| *UAS-snmase*<sup>RNAi</sup> | Unaffected | Increased number and size | Increased number and reduced size | Slightly larger in size |
| *UAS-sply*<sup>RNAi</sup> | Unaffected | Unaffected | Unaffected | Unaffected |
| *UAS-sk1*<sup>RNAi</sup> | Increased number | Unaffected | Unaffected | Unaffected |
| *UAS-sk2*<sup>RNAi</sup> | Unaffected | Unaffected | Unaffected | Unaffected |
| *UAS-ksr*<sup>RNAi</sup> | Reduced number | Reduced number | Reduced number | Reduced number |
| *UAS-cpt1*<sup>RNAi</sup> | Increased number and size | Increased number and size | No change | Reduced number |
| *UAS-fabp1*<sup>RNAi</sup> | Unaffected | Unaffected | Unaffected | Unaffected |
| *UAS-fatp1*<sup>RNAi</sup> | Sporadic increase in size | Unaffected | Unaffected | Unaffected |

**Table 2:**
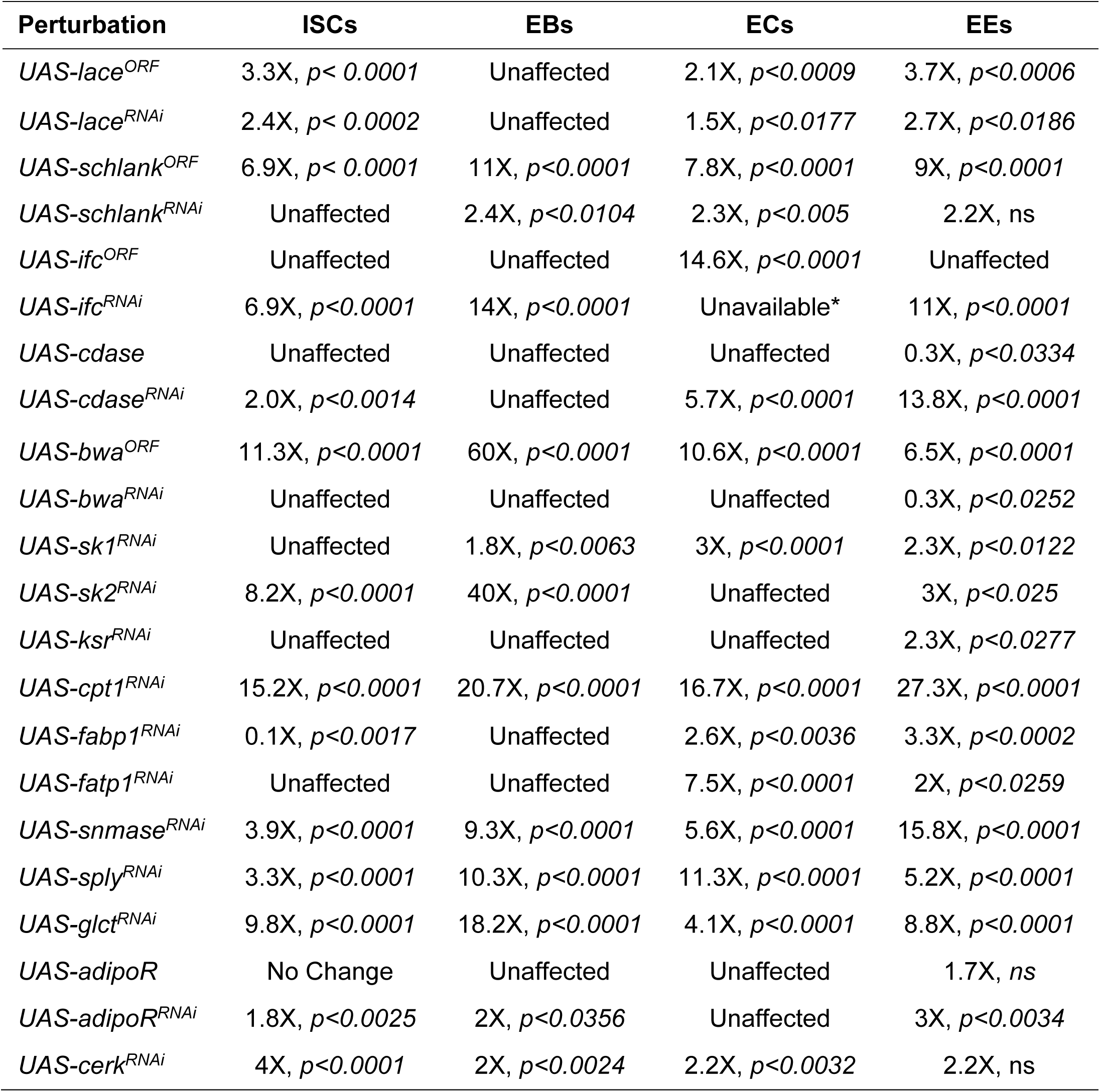
Cell-autonomous and non-cell-autonomous effects of changes in the levels of ceramide metabolic enzymes. Shown here is a summary of fold change in median ISC proliferation relative to control in each condition with statistical significance. The fold change between 0.5 to 1.5 with p>0.05 was treated as ‘unaffected’ in that condition. * The cross-setup did not generate progeny with the right genotype.

| **Perturbation** | **ISCs** | **EBs** | **ECs** | **EEs** |
| --- | --- | --- | --- | --- |
| *UAS-lace<sup>ORF</sup>* | 3.3X, $p < 0.0001$ | Unaffected | 2.1X, $p < 0.0009$ | 3.7X, $p < 0.0006$ |
| *UAS-lace<sup>RNAi</sup>* | 2.4X, $p < 0.0002$ | Unaffected | 1.5X, $p < 0.0177$ | 2.7X, $p < 0.0186$ |
| *UAS-schlank<sup>ORF</sup>* | 6.9X, $p < 0.0001$ | 11X, $p < 0.0001$ | 7.8X, $p < 0.0001$ | 9X, $p < 0.0001$ |
| *UAS-schlank<sup>RNAi</sup>* | Unaffected | 2.4X, $p < 0.0104$ | 2.3X, $p < 0.005$ | 2.2X, ns |
| *UAS-ifc<sup>ORF</sup>* | Unaffected | Unaffected | 14.6X, $p < 0.0001$ | Unaffected |
| *UAS-ifc<sup>RNAi</sup>* | 6.9X, $p < 0.0001$ | 14X, $p < 0.0001$ | Unavailable* | 11X, $p < 0.0001$ |
| *UAS-cdase* | Unaffected | Unaffected | Unaffected | 0.3X, $p < 0.0334$ |
| *UAS-cdase<sup>RNAi</sup>* | 2.0X, $p < 0.0014$ | Unaffected | 5.7X, $p < 0.0001$ | 13.8X, $p < 0.0001$ |
| *UAS-bwa<sup>ORF</sup>* | 11.3X, $p < 0.0001$ | 60X, $p < 0.0001$ | 10.6X, $p < 0.0001$ | 6.5X, $p < 0.0001$ |
| *UAS-bwa<sup>RNAi</sup>* | Unaffected | Unaffected | Unaffected | 0.3X, $p < 0.0252$ |
| *UAS-sk1<sup>RNAi</sup>* | Unaffected | 1.8X, $p < 0.0063$ | 3X, $p < 0.0001$ | 2.3X, $p < 0.0122$ |
| *UAS-sk2<sup>RNAi</sup>* | 8.2X, $p < 0.0001$ | 40X, $p < 0.0001$ | Unaffected | 3X, $p < 0.025$ |
| *UAS-ksr<sup>RNAi</sup>* | Unaffected | Unaffected | Unaffected | 2.3X, $p < 0.0277$ |
| *UAS-cpt1<sup>RNAi</sup>* | 15.2X, $p < 0.0001$ | 20.7X, $p < 0.0001$ | 16.7X, $p < 0.0001$ | 27.3X, $p < 0.0001$ |
| *UAS-fabp1<sup>RNAi</sup>* | 0.1X, $p < 0.0017$ | Unaffected | 2.6X, $p < 0.0036$ | 3.3X, $p < 0.0002$ |
| *UAS-fatp1<sup>RNAi</sup>* | Unaffected | Unaffected | 7.5X, $p < 0.0001$ | 2X, $p < 0.0259$ |
| *UAS-snmase<sup>RNAi</sup>* | 3.9X, $p < 0.0001$ | 9.3X, $p < 0.0001$ | 5.6X, $p < 0.0001$ | 15.8X, $p < 0.0001$ |
| *UAS-sply<sup>RNAi</sup>* | 3.3X, $p < 0.0001$ | 10.3X, $p < 0.0001$ | 11.3X, $p < 0.0001$ | 5.2X, $p < 0.0001$ |
| *UAS-glct<sup>RNAi</sup>* | 9.8X, $p < 0.0001$ | 18.2X, $p < 0.0001$ | 4.1X, $p < 0.0001$ | 8.8X, $p < 0.0001$ |
| *UAS-adipoR* | No Change | Unaffected | Unaffected | 1.7X, ns |
| *UAS-adipoR<sup>RNAi</sup>* | 1.8X, $p < 0.0025$ | 2X, $p < 0.0356$ | Unaffected | 3X, $p < 0.0034$ |
| *UAS-cerk<sup>RNAi</sup>* | 4X, $p < 0.0001$ | 2X, $p < 0.0024$ | 2.2X, $p < 0.0032$ | 2.2X, ns |

### Brainwashing (bwa) promotes ISC proliferation autonomously and non-cell autonomously

Of the genes we tested, the predicted alkaline ceramidase named *brain washing* (bwa) gave some of the strongest effects. Bwa shares significant amino acid homology with the human alkaline ceramidase Acer2. However, unlike ACER2, Bwa has been shown to have no ceramidase activity^22^, Figure 3B, and the specific nature of its true enzymatic activity is unknown. Bwa genetically interacts with the other predicted *Drosophila* ceramidase (*Cdase*) and with ceramide kinase (*cerk*), and can regulate sphingolipid metabolism. To understand the role of bwa in the intestine, we altered the levels of bwa in each of the different intestinal cell types as described above. Bwa gene function was either elevated or reduced at the mRNA transcript level for five days. Compared to controls, bwa gain-of-function (GoF) increased the number and size of ISCs and EBs while no changes were visible for ECs and EEs (Figures 1B, H). The knockdown of Bwa was associated with a small increase in the number and size of ISCs and EBs, but there was no change in the number or size of the ECs and EEs (Figure 1B lower panel). *Bwa* overexpression in any cell type increased ISC proliferation, indicating cell-autonomous and non-cell-autonomous effects (Figure 1C). The proliferation phenotype of ISCs was strongest when *bwa* was overexpressed in EBs, which gave an average of ∼250 PH3^+^ mitotic cells per midgut. In contrast, expression in other cell types usually gave ∼100 PH3^+^ cells per midgut compared to the normal ∼10 PH3^+^ cells. The knockdown of *bwa* was associated with a small but significant increase in ISC proliferation in the midgut. However, heterozygous *bwa* mutants showed no difference in ISC proliferation (Supplementary Figure 4), suggesting that the robust ISC proliferation we saw in the *bwa* GoF condition, and the small increase in ISC proliferation in *bwa* knockdown conditions are two distinct phenotypes.

Since increasing *bwa* in EBs lead to a large increase in ISCs proliferation, we sought to assess whether *bwa* levels could affect the differentiation potential of ISCs in the same condition. Compared to controls, the number of both ISCs and EBs increased in the *bwa* gain of function in EB conditions, while no significant change in the number of EEs was observed when bwa was overexpressed in EBs. (Figure 1D-G). On the other hand, *bwa* knockdown in EBs increased the number of EEs, compared to controls, without significantly changing the numbers of ISCs or EBs (Figure 1D-G). There was an increased activity of the Notch signaling pathway when bwa was increased in the progenitor cells (ISCs +EBs), however, no change in the relative number of ISCs or EBs was observed (Supplementary Figure 2). These results suggest that bwa activity has no strong effects on cell type specification or differentiation in the ISC lineage but instead has a strong non-cell autonomous effect on ISC proliferation.

### Bwa-mediated ISC proliferation requires ceramide synthesis but not Sphingosine-1-Phosphate (S1P)

Sphingosine-1-phosphate (S1P) is a known mitogenic factor identified in several studies^23^. As shown in Figure 1A, S1P is a product of Sphingosine phosphorylation by the isoenzymes-sphingosine kinase 1 and 2 (Sk1, Sk2). To test whether the bwa GoF-mediated ISC proliferation depends upon increased levels of S1P, we knocked down *Sk1* and/or *Sk2* in the *bwa* GoF condition. In all three conditions, *Sk* knockdown had no repressive effect on ISC proliferation induced by over-expressed *bwa* (Figure 2A). The opposite was observed: the double knockdown of *Sk1* and *Sk2* further increased ISC proliferation driven by bwa over-expression; These results suggest that bwa GoF-induced ISC proliferation is independent of the S1P activity.

**Figure 2.**
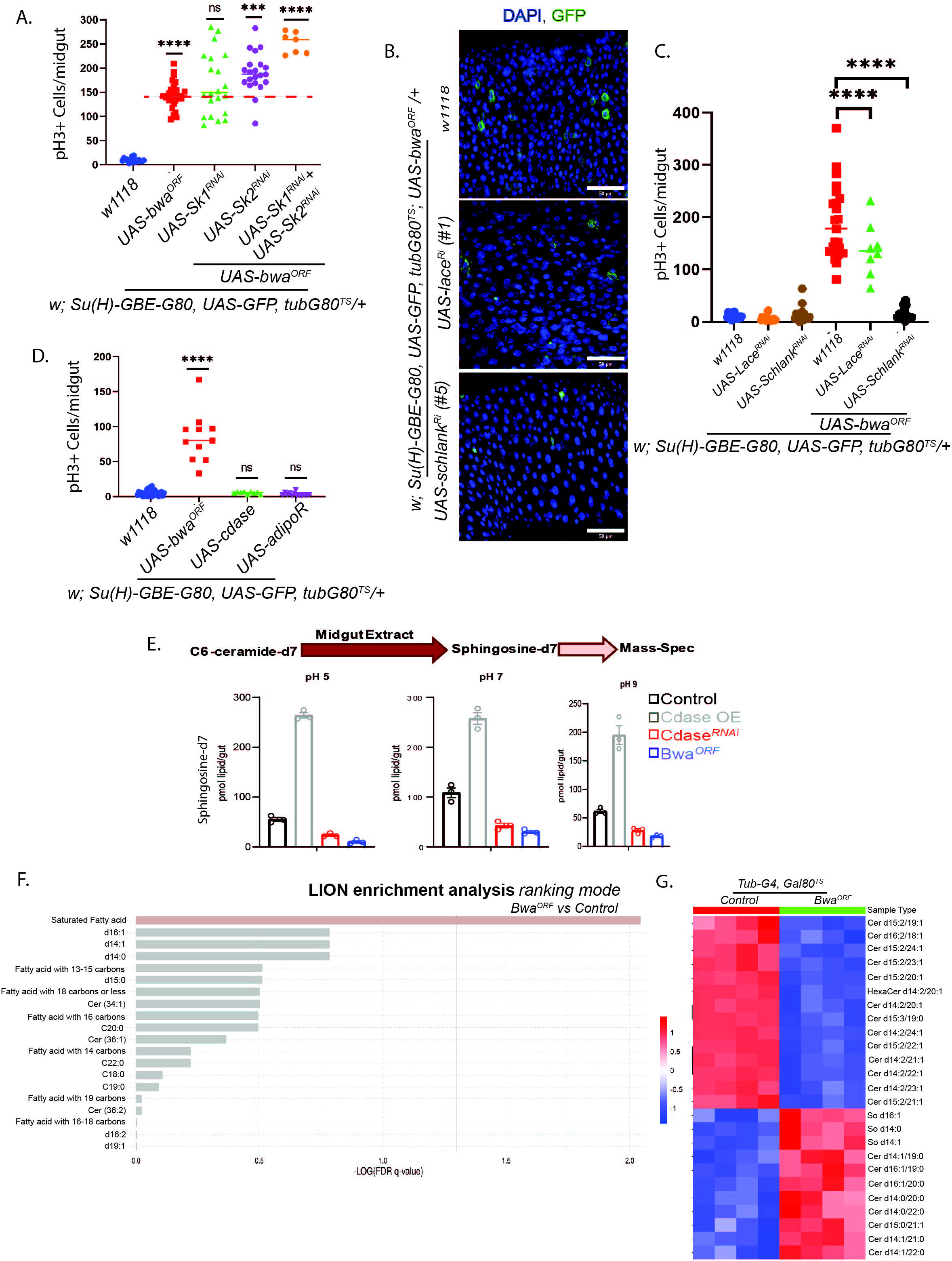
Bwa Gain of function-induced ISC proliferation is independent of Sphingosine-1-Phosphate (S1P) or ceramidase activity. A. Graph showing knock down (KD) of sphingosine kinases 1 and 2 do not rescue the Bwa Gain of function-induced excessive ISC proliferation phenotype either as single knockdown or double knockdown. B. Images showing EB-specific Bwa Gain of function leads to cell growth that is rescued by lace or schlank knockdown. C. Graph showing cell non-autonomous bwa Gain of function induced ISC proliferation phenotype is rescued by lace or schlank knockdown. D. EB-specific Bwa Gain of function-induced ISC proliferation is not phenocopied by other known ceramidases such as ceramidase (Cdase), or adiponectin receptor (adipoR representing all the different receptor sub-types). E. Graphs showing ceramidase activity of the midgut extracts from control, Gain of function/Loss-of-function ceramidase, and Bwa Gain of function genotypes as measured through Mass-spec and normalized to the number of midguts. F. Graph showing enrichment ranking of different fatty acids in untargeted lipidomic (LC-MS) analysis performed using LION enrichment software (https://www.lipidmaps.org/resources/tools/stats). G. Heatmap showing statistical enrichment of top sphingolipid species showing maximum differences in control versus Bwa Gain of function conditions. * represents values e.g. *\* p< 0.05 and, ** p< 0.005, *** p< 0.0005, **** p< 0.00005*.

Since bwa GoF-induced ISC proliferation is independent of S1P activation, we assessed whether the phenotype is even a sphingolipid-dependent event. We knocked down components of the early stages in the ceramide synthetic pathway such as *lace* and *schlank* using RNAi constructs both in wild type and in bwa GoF conditions in the EBs. While knockdown of lace and schlank in the wild type EBs did not affect ISC proliferation, this knockdown severely reduced ISC proliferation in bwa of conditions (Figure 2B, C), suggesting that bwa GoF induced ISC proliferation is a sphingolipid-dependent phenomenon.

### Bwa is not a ceramidase but promotes the accumulation of saturated sphingolipids and free fatty acids

Bwa is a putative alkaline ceramidase and shows 46% amino acid identity and 70% similarity to the human alkaline ceramidase gene 1^22^. Phylogenetic analysis shows that Bwa is closer to human Acer2^24^. However, no actual ceramidase activity was found for bwa in head extracts of flies^22^, raising the question of whether bwa encodes an active ceramidase enzyme. Rather, genes involved in oxidative stress pathways were upregulated in *bwa* homozygous mutants^25^. In our tests, however, over-expression of c*atalase-A* (Cat-A) and *superoxide dismutase 1* (SOD1) together with *bwa^ORF^* did not affect the *bwa* GoF-induced ISC proliferation (Figure 3G), suggesting that bwa GoF-induced phenotype has little if any connection to oxidative stress.

**Figure 3.**
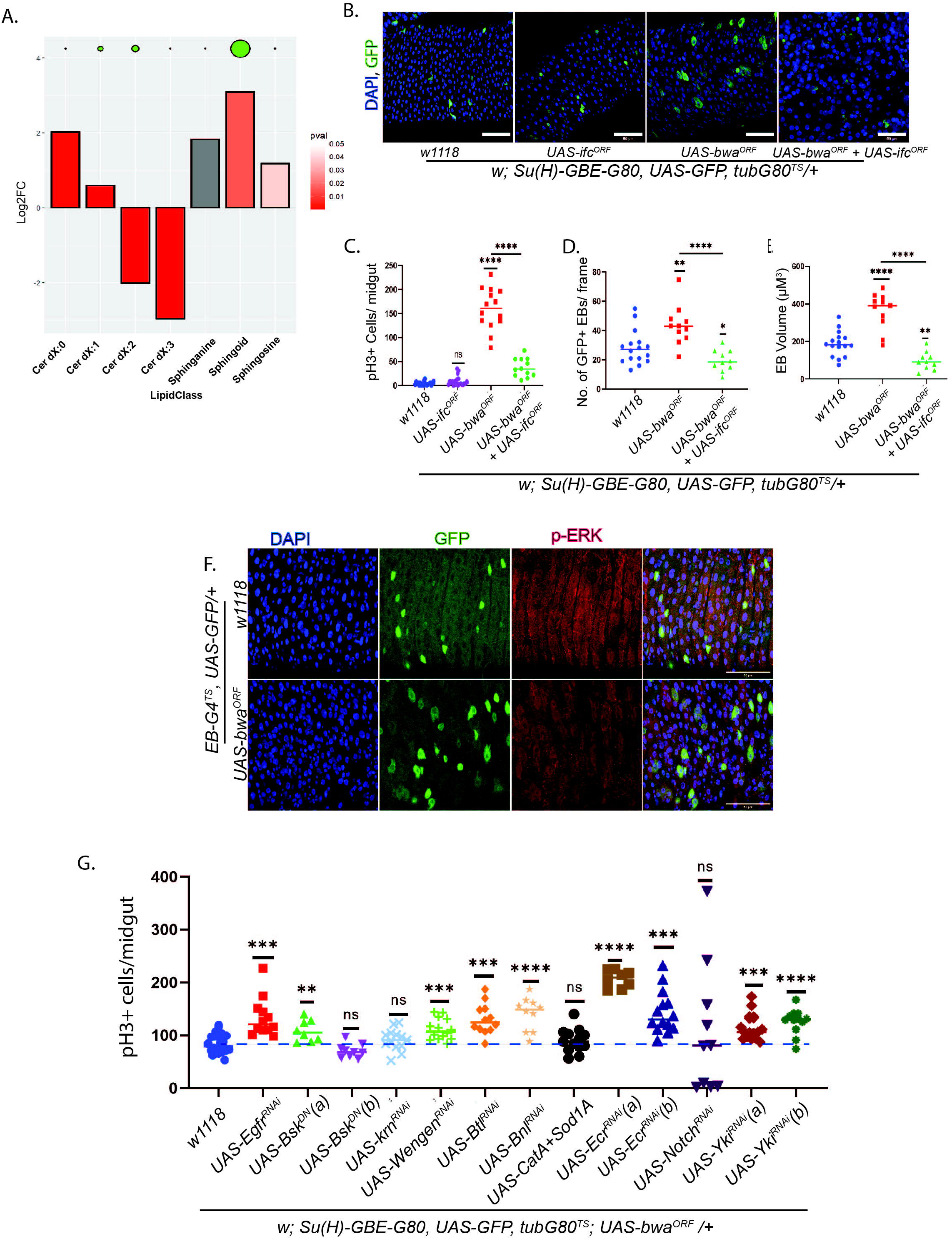
Bwa over-expression is associated with the accumulation of saturated sphingolipids and free fatty acids, promoting ISC proliferation. A. Bar graph showing class enrichment analysis of the different sphingolipid classes. B-E. Images showing EBs (green) and graphs showing the proportion of proliferating ISCs, EBs, and EB size (volume) in Bwa Gain of function alone and Bwa with Ifc Gain of function conditions and the respective controls. F. Images show no difference in p-ERK activity in EB-specific control versus Bwa gain in function conditions. G. Pseudo-epistasis test graph shows usual damage responsive pathways such as EGFR, JNK, FGF, ECR, Notch, Hippo or antioxidant pathways do not affect Bwa gain of function-induced ISC proliferation. Scale bars represent 50 µM, * represents values e.g. *\* p< 0.05 and, ** p< 0.005, *** p< 0.0005, **** p< 0.00005*.

We tested whether known ceramidases could phenocopy the bwa effect in the midgut. EB-specific over-expression of the pan ceramidase, *Cdase,* or the adiponectin receptor (*adipoR),* which as been shown to exhibit ceramidase activity in mammals^26^ did not promote any ISC proliferation compared to the controls (Figure 2D), suggesting *bwa* does not increase ISC proliferation via ceramidase activity. To more directly assess the potential ceramidase activity of *bwa* in the midgut, we performed a ceramidase assay using midgut extracts from four genotypes corresponding to *w^1118^* (control), *Cdase* overexpression, *Cdase* knockdown, and *bwa* overexpression. Deuterated C6-ceramide-d7 was incubated at three different pH conditions with the extracts, and the reaction product was run through mass-spectroscopic analyses to assess the levels of sphingosine-d7. In all three pH conditions. *Cdase* overexpression caused an approximately three-fold increase in the levels of Sphingosine-d7, whereas both *Cdase* and *bwa* overexpression showed reduced levels of sphingosine-d7 compared to the *w^1118^* control group. These results confirm the results of Yuan et al.^22^ in suggesting that bwa does not encode an active ceramidase (Figure 2E). Indeed, our results in Figure 2E indicate that *bwa* overexpression might have some dominant-negative activity toward other ceramidases.

Given these results, we sought to assess the overall effects of *bwa* on the lipid composition of intestinal cells. We performed a high-precision targeted and untargeted lipidomic study using liquid- and gas-chromatography and mass spectroscopy (LC/MS and GC/MS) on the midgut extracts from control, *bwa* overexpressing, and *bwa*-RNAi knockdown genotypes. Enrichment analysis of untargeted lipidomics data showed that fully saturated and mono-unsaturated fatty acids were the most significantly enriched fatty acid in midguts with *<u>bwa</u>* overexpression, compared to control midguts, while no significant changes were observed in the *bwa* knockdown condition (Figure 2F, Supplementary figure 3). Targeted analysis of the sphingolipid species involved in the ceramide metabolism showed a similar up-regulation of those with saturated or mono-unsaturated sphingoid and acyl chains in the *bwa* GoF condition (Figure 2G). On the other hand, unsaturated sphingolipids were severely downregulated in the *bwa* of condition (Figure 2G). Similarly, ceramides with two or three double bonds in the sphingoid and acyl chains were downregulated by *bwa* overexpression. In contrast, those with zero or only one double bond were not downregulated (Figure 3A). These results suggest that *bwa* overexpression can increase the saturation levels of sphingolipids and free fatty acids.

To test this hypothesis, we performed an experiment in which we fed the flies Myristic Acid, a saturated fatty acid, or Linoleic acid, an unsaturated fatty acid, as well as appropriate controls. After five days, we found that flies feeding on Myristic acid had significantly higher ISC proliferation than vehicle controls or those feeding on Linoleic acid (Supplementary Figure 5). Although the effect was much milder than seen with bwa overexpression, this result is consistent with the notion that lipid saturation can stimulate ISC proliferation.

Following this line of investigation, we hypothesized that if *bwa*-dependent enrichment of saturated lipids is causal for *bwa*-induced ISC proliferation, then reversing this enrichment using enzymes that drive lipid de-saturation should be able to suppress *bwa*-dependent ISC proliferation. Hence, we co-expressed a ceramide desaturase enzyme - *infertile crescent (ifc)* – along with *bwa*. The *ifc* gene encodes a homolog of human dihydroceramide desaturase 1 (*Des1*), which has been shown to introduce a double bond on dihydroceramide to convert it to ceramide^27^. When expressed alone in EBs, *ifc* did not affect EB size or number, or the amount of ISC proliferation (Figure 3B-E). However, EB-directed co-expression of *ifc^ORF^* and *bwa^ORF^*led to a reduction in the number and size of EBs and the proportion of proliferating ISCs that would typically result from *bwa^ORF^* expression alone (Figure 3B-E). These complementary lipidomic, morphological, and immunofluorescence imaging and image quantification data sets suggest that the saturation status of the sphingolipids is instructive for ISC proliferation in the *bwa* overexpression condition.

### Bwa activates p38^MAPK^ and Upd/JAK/Stat inflammatory signaling to promote ISC proliferation

As the activation ISC proliferation was strongest when *bwa* was overexpressed in ISC-adjacent enteroblasts (EBs), we hypothesized that an excess of saturated sphingolipids and or free fatty acids might send a paracrine signal to ISCs to trigger their division. To test this potential non-cell autonomous mechanism, we investigated several signaling pathways that are known to promote ISC activation. Immunofluorescence staining of *bwa*- overexpressing midguts showed no increase in p-ERK, suggesting that signaling via the EGFR/RAS/RAF/ERK was not a key determinant (Figure 3F). In addition, epistasis tests with components of general stress response pathways, including EGFR, JNK, Wnt, Fgf, EcR, Notch, Hippo, and oxidative stress signaling pathways, showed that *bwa*-induced ISC proliferation could not be suppressed by depletion of components in any of these pathways (Figure 3G). These results suggested that the *bwa*-mediated induction of ISC proliferation is not a general damage response phenotype.

Besides damage response pathways, inflammation is a potent regulator of ISC proliferation^28^. Excess sphingolipids, long-chain fatty acids, and high-fat diets have been shown to stimulate inflammatory signaling in other contexts^1,29,30^. To determine whether saturated fatty acid accumulation might be able to trigger an inflammatory response in the fly gut and thus promote ISC proliferation, we performed experiments targeting the P38-Upd/Jak/Stat axis, an inflammatory regeneration response pathway in flies^31,32^. Immunofluorescence imaging showed that both phospho-P38, the active form of a kinase that can induce inflammatory cytokines, and Upd3, an inflammatory cytokine, were both up-regulated in the midgut by bwa-overexpression (Figure 4A, B, Supplementary Figure 6). Consistent with previous reports that Upd3 is mitogenic in the gut^33,34^, overexpression of Upd3 in EBs strongly increased ISC proliferation, and also increased the number of Dl^+^ ISC and (Figure 4C, D). We also performed complementary epistasis tests in which Upd2 or Upd3 was depleted by RNAi in EBs that co-expressed *bwa*. These tests showed that Upd2- or Upd3-RNAi was able to reduce the increase in EB size and numbers caused by overexpressed *bwa*, and also reduce the *bwa*-driven increase in ISC proliferation (Figure 4E, F). These results support our proposition that *bwa* over-expression causes an accumulation of saturated sphingolipids and other lipids, which in turn triggers P38/Upd-dependent inflammatory signaling that promotes EB cell growth and (non-cell autonomously) ISC proliferation (Figure 4G).

**Figure 4.**
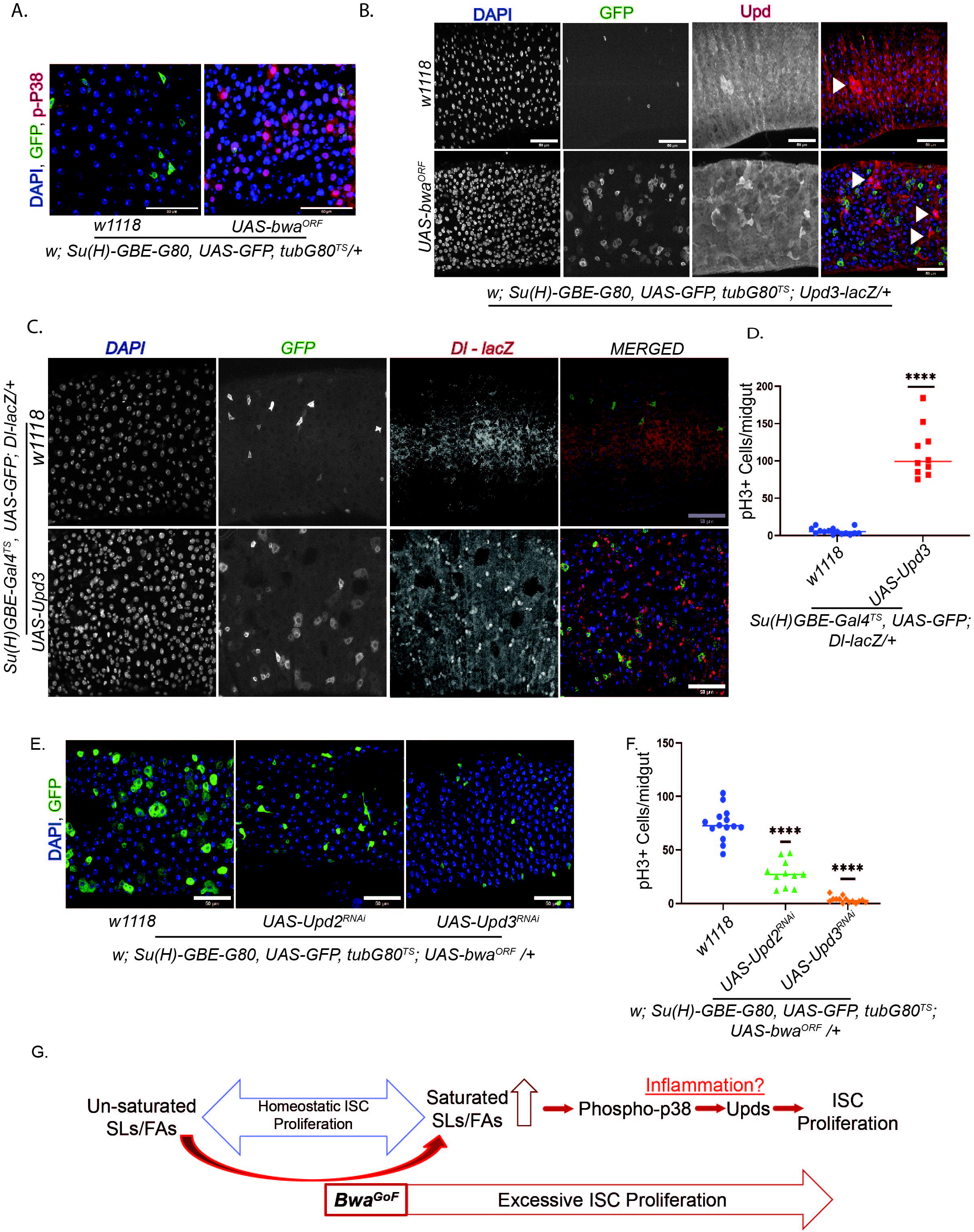
Bwa over-expression-induced accumulation of saturated sphingolipids activates inflammatory signals such as p38 and Upds to promote ISC proliferation. A. Images showing pro-inflammatory phosphor-P38 activation (red) in Bwa Gain of function function conditions. B. Images showing inflammatory cytokine Upd being upregulated in the midgut with Bwa Gain of function condition compared to the controls. C. Images showing EB sizes and numbers in control versus Upd Gain of function conditions. D. Graph showing cell non-autonomous ISC proliferation in the EB-specific Upd3 Gain of function condition compared to the controls. E. Images showing the interaction between Bwa Gain of function and Upd2 and Upd3 knockdown conditions. F. Graphs showing ISC proliferation in EB-specific Bwa Gain of function coupled with knockdown of Upd2 and Upd3. G. Cartoon showing the relationship between saturated versus unsaturated fatty acids and stem cell proliferation and the role of Bwa in tilting the equilibrium wherein Bwa-mediated enrichment of saturated sphingolipids or fatty acids activate components of the inflammatory pathway such as phospho-P38 and Upds, which in turn prompt excessive ISC proliferation. Scale bars represent 50 µM, * represents values e.g. *\* p< 0.05 and, ** p< 0.005, *** p< 0.0005, **** p< 0.00005*.

## DISCUSSION

We found that cell-type-specific genetic alterations of the ceramide metabolic enzymes have robust cell-autonomous and non-cell-autonomous effects on cell morphologies and behaviors in the *Drosophila* midgut (Supplementary Figure 1, Supplementary Figure 2, Tables 1 and 2). We have shown that *bwa* over-expression is associated with higher Notch activity in the ISC lineage and an increase in the numbers of all the cell types (Supplementary Figure 2, and Figures 1B, C), suggesting that *bwa* gain of function increased ISC proliferation without affecting the differentiation potential of the EBs. Targeted and untargeted lipidomic analysis showed that *bwa* overexpression caused a robust accumulation of saturated sphingolipid metabolites and saturated fatty acids (Figure 3C, D, E). In addition, co-expression of the gene *infertile crescent (ifc),* which desaturates dihydroceramide by introducing a carbon chain double bond, counteracted the pro-proliferative effect of bwa overexpression (Figure 3F, G). Furthermore, pro-inflammatory interleukins including Upd2 and Upd3 are necessary for *bwa*-induced ISC proliferation (Figure 4E, F). Together, these results suggest that bwa gain of function increases the levels of saturated sphingolipids and possibly other saturated lipids, which leads to the activation of inflammatory signaling to promote ISC proliferation. We propose that the saturation status of the sphingolipids is a key determinant of ISC proliferation, which is associated with GI homeostasis, as summarized in Figure 4G.

The role of lipid metabolism in affecting intestinal stem cell physiology is appreciated^35,36^, though the underlying mechanisms are not well understood. Recent studies have established that high-fat diets containing significant amounts of sphingolipids promote ISC proliferation^37^. Ceramide has recently been shown to promote stemness and increase the risk of tumorigenesis^11^. At the same time, earlier studies have established the relative amount of S1P, a ceramide metabolite, to ceramide in the tumor versus the serum may alter the rate of tumor growth and/or progression^38^. The role of ceramide and its metabolites, thus, encompasses many aspects of gastrointestinal biology-from maintaining gut homeostasis to promoting diseases such as colorectal cancer, and inflammatory bowel diseases^39,40,41,42^. We can posit that different metabolites in the ceramide pathway (Figure 1A) may trigger different cellular signaling nodes to effect distinct responses in cellular physiology. In addition, each intestinal cell type, having its own set of intracellular signaling networks, may respond differently to the signaling cues affected by the ceramide metabolites. Depending on which network is dominantly affected, alterations in ceramide metabolites may change diverse physiological parameters, such as cell death, differentiation, proliferation, or senescence, and also produce non-cell-autonomous signaling factors. However, few if any studies have been carried out that reveal a definitive answer for how a single cell type of any type might respond to specific alterations in ceramide metabolism. Our work as presented here addresses some of these issues by offering a framework for understanding how intestinal stem cells and their lineages respond when ceramide metabolites are altered, either cell-autonomously or non-cell-autonomously.

In this study, by altering the activity of each ceramide metabolism gene in each gut cell type, we uncovered many instances of ceramide metabolism alterations that can perturb gut homeostasis. Some of these effects were cell autonomous, and others were non-cell autonomous. A thorough analysis of each perturbation and its effect has not been possible to pursue. Nor have we established whether *bwa* acts as an enzyme or a regulatory protein that alters the action of other enzymes in the ceramide/sphingolipid metabolic pathway. However, *bwa* does not appear to act as a simple ceramidase like its human homolog, the alkaline ceramidase ACER2. This raises the tantalizing possibility that Acer2 might also target unsaturated long-chain sphingolipids and promote their hydrolysis to saturated sphingolipids. While this study has produced many phenotypes and questions, answering all of them was beyond the scope the work we present here. However, by studying the phenotype produced by *bwa* expression in EBs, and its impact on ISCs, we have been able to decipher a general biological aspect of sphingolipids, namely that saturated sphingolipids can promote cell growth and inflammatory signaling that foments the ectopic proliferation of ISCs (Figure 4G). These findings offer clues to not only how ceramides and other sphingolipid metabolites can affect a given cell type cell-autonomously or non-cell-autonomously but also shed light on how the saturation status of the long-chain fatty acids might affect gastrointestinal health.

## MATERIALS AND METHODS

### Fly stocks and maintenance

Drosophila stocks were maintained at 180C on a 12:12 hour light: dark cycle on standard fly food containing propionic acid (Sigma) and Tegosept (Genesee). The following Gal4 lines were used for tissue and cell-type specific expression: ISC-G4ts (yw; EsgG4, UAS GFP, tubG80ts/Cyo; Su(H)-GBE-Gal80/TM6B), EB-G4ts (yw; Su(H)-GBE-Gal4, UAS GFP, tubGal80ts/Cyo), EC-G4ts (w; Myo1A-Gal4, tubGal80ts, UASGFP/Cyo), EE-G4ts (w; ProsV1Gal4, UASGFP/Cyo; tubGal80ts/TM6B^)20,21.^ The following UAS strains were used: *UAS-lace^RNAi^*(VDRC# v110181), *UAS-Schlank^RNAi^* (BDSC# 29340), *UAS-ifc^RNAi^*(BDSC# 32514), *UAS-ifc^RNAi^* (VDRC# 106665), *UAS-Cdase^RNAi^*(BDSC# 36764), *UAS-bwa^RNAi^* (BDSC# 36765), *UAS-Sk1^RNAi^*(VDRC# v30054), *UAS-Sk2^RNAi^* (BDSC# 35570), *UAS-Sply^RNAi^*(BDSC# 42834), *UAS-Sply^RNAi^* (BDSC# 43304), *UAS-Sply^RNAi^*(VDRC# v103485), *UAS-SnMase^RNAi^* (BDSC# 42834), *UAS-fatp1^RNAi^*(BDSC# 50709), *UAS-Cpt1(whd)^RNAi^* (BDSC# 33635), *UAS-Cpt1(whd)^RNAi^* (BDSC# 34066), *UAS-Cpt1(whd)^RNAi^*(VDRC# v105400), *UAS-fabp1^RNAi^* (BDSC# 34685), *UAS-Ksr^RNAi^* (BDSC# 32937), *UAS-Schlank^ORF^* (F003905), *UAS-lace^ORF^* (F003905), *UAS-ifc^ORF^* (F003887), *UAS-bwa^ORF^* (F004012).

These genotypes are not used in the powerpoint, but are still the same genotype (different lines): *UAS-Sk1^RNAi^* (BDSC# 36747), *UAS-Sk2^RNAi^* (BDSC#36741), *UAS-Sk2^RNAi^* (VDRC# v101018), *UAS-bwa^RNAi^*(BDSC# 29409), *UAS-bwa^RNAi^* (VDRC# v101366), *UAS-Ras^V^*^12^ (BDSC# 4847) and *UAS-Notch^DN^* (92), *UAS-Notch^RNAi^* (BDSC# 7077), *UAS-lace^RNAi^* (BDSC# 51475), *UAS-fatp1^RNAi^* (BDSC# 55273), *UAS-fatp1^RNAi^* (VDRC# v100124), *UAS-Ksr^RNAi^* (VDRC# v110621). The following mutants and reporter lines were used: *w1118* (BDSC# 3605), esg-lacZ606 (BDSC# 10359), Dl-lacZ05151 (BDSC# 11651) and Su(H)-lacZ (BDSC# 83352). *Bwa^e02081^* and *Bwa^KG0406S^* lines were used as the bwa heterozygous mutants. For crosses involving driver lines with “tubulin-Gal4, Gal80ts” were set up at 180C, F1 adult progeny were collected within two days of emerging from the pupae. These flies were transferred to 29^0^C for the transgene expression including *RNAi* construct of interest. Flies were used within ten days of emerging for all the stress experiments. Flies were transferred to new food every two days except for weekends

### Dissections and Immunohistochemistry

Adult female flies were selected and dissected for the midgut, including the proventriculus and the hindgut. Following dissections on in 1x PBS (130mM NaCl, 70mM Na2HPO4, 30mM NaH2PO4), the midguts were fixed in 4% Paraformaldehyde (Ted Pella Inc.) in 1x PBS for 30 minutes at room temperature. After 3 rinses for 5 min each with 1x PBS-T (1x PBS, 0.2% TritonX-100), the tissues were incubated in blocking PBT solution (1x PBS, 2% BSA, 4% Goat Serum, 0.2% TritonX-100) at room temperature for 45 minutes. The primary antibody was added in PBT by incubation for 16 hours at 4°C. The primary antibody was washed 3 times for 5 min each at room temperature with PBS-T. Secondary antibodies and DAPI (Sigma) are added for 2 hours at room temperature. Excess antibodies are washed in PBS-T (as before) and the midguts are mounted in Vectashield (Vector). Primary antibodies: rabbit-anti-pH3 (Millipore 1:1000) (mitosis marker), mouse-anti-Prospero (DSHB 1:200), rabbit-anti-β-gal (origin unknown 1:200). Secondary antibodies: DAPI (Sigma, 1:2000 of 10mg/ml stock) (staining of the nuclei), against mouse or rabbit conjugated to Alexa Fluor 568 (Invitrogen cat. No. A-11011) were used at 1:500.

### PH3 Staining, counting, statistical analysis, and graphical representation

Once mounted, the numbers of pH3-positive cells were counted under the fluorescent microscope (Nikon Eclipse Ti) at 20x magnification along the whole midgut. Each dot indicates the PH3 count for a single midgut. Graphs were generated using GraphPad Prism 9, San Diego, CA. Statistical significance was assessed using an unpaired two-tailed Mann-Whitney t-test. The comparisons were between control and test samples unless otherwise indicated.

### Microscopy and Image Quantification

The numbers of PH3-positive cells were counted under the fluorescent microscope Nikon Eclipse Ti at 20x magnification along the whole midgut. Graphs were generated using Graphpad Prism 8-10 by calculating the average and standard deviation values. Midgut sectional images were acquired with a Leica SP8 Confocal microscope, equipped with GFP, DAPI, and Alexa Fluor 568 filters. Stacks of optical sections were acquired using this microscope with a 40x water objective (zoom 1.28x, line average 1, line accumulation 3, speed 700). Images were processed using ImageJ for midgut visualization. Images were uploaded into ImageJ via a *.lif file, and color mode made composite. Z project stacks were selected, and max intensity was implemented.

Once processed, confocal images were quantified using the IMARIS software (version 10.0, Oxford Instruments). The images were uploaded onto the software and converted to .ims files from .lif files. Image processing was performed by reducing the background and baseline noise of over-saturated channels (view image proc). Surfaces were generated to quantify each channel (DAPI, GFP, and RFP, corresponding to blue, green, and red). The DAPI signals were quantified by adjusting the intensity threshold to select all possible nuclei, followed by filtering based on the number of voxels and the sphericity of the signals. The GFP signals were quantified by adjusting the intensity threshold and using the quality setting to remove lower-quality signals. Like DAPI, the RFP signals were quantified by adjusting the intensity threshold, followed by filtering based on the number of voxels and sphericity of the selected signals. Once quantified, the signal’s volume and area statistics were saved for each channel as a ‘*.xlsx’ file. The number of signals for each channel was determined, as well as the corresponding means and standard deviations, using Microsoft Excel. Graphs and statistical analyses were generated using GraphPad Prism 9.

### Ceramidase Assay

A total of 10 midguts per replicate per genotype were dissected out and collected in 100 µl of ice-cold non-denaturing lysis buffer (20 mM Tris-HCl pH 7.5, 150 mM NaCl, 1 mM Na2EDTA, 1 mM EGTA, 1% Triton, 2.5 mM sodium pyrophosphate, 1 mM beta-glycerophosphate, 1mM sodium orthovanadate with 1x Complete Protease Inhibitor solution). To prepare extracts, the midguts were ground with a green pestle and mixed gently with an additional 1200 µl of ice-cold non-denaturing lysis buffer, spun at maximum speed at 4^0^C, and the clear supernatant (∼500 µl) was transferred to ice-cold Eppendorf tubes. To assess the ceramidase activity, the midgut extracts were incubated with C6-ceramides-d7, and the resulting sphingosine-d7 (the resulting product of ceramidase-mediated degradation of C6-ceramide-d7) levels were monitored.

### Lipidomics

For lipidomic experiments, a total of 40 midguts per replicate was collected in ice-cold 1xPBS, transferred to 2.0 mL ceramic bead mill tubes (Qiagen Catalog Number 13116-50) and centrifuged at maximum speed. The PBS was then carefully removed and the samples were frozen in -80^0^C. Sample extraction protocol for lipidomics based on Matyash et al. J Lipid Res 49(5) (2008) 1137-1146 and was processed as described previously^43^.

### Fatty acid feeding

Saturated myristic acid (Sigma-Aldrich, M3128) and unsaturated linoleic acid (Sigma-Aldrich, L1376) were dissolved in 200 proof ethanol (Decon labs Inc., SKU: 07-678-003) to make a 50mM solution of the fatty acids. For each fatty acid solution, a yeast paste was prepared by mixing 2 gms of yeast powder with 2.5 mls of water. Next, 1 ml of the fatty acid solution was mixed thoroughly with the yeast paste. To make the yeast paste firmer with fatty acids and to avoid flies being stuck to the paste, a further 0.5 gms of yeast powder was added again. For controls, an equal volume (1ml) of 200-proof ethanol was mixed with yeast paste and powder. These yeast pastes were added to the vials in equal amounts to feed a day-old w1118 flies for three days. After three days of feeding, the flies were dissected and midguts were stained for pH3 as previously described.

## ACKNOWLEDGEMENTS

We are grateful to the University of Utah Cell Imaging and Metabolomics Cores for their expert assistance. This work was supported by NIH grants DK131609 to SAS and GM140900 to BAE, and Huntsman Cancer Institute grant 23-TSA-02.

