## Supplementary figures and images for "Bwa, an ortholog of alkaline ceramidase-ACER2, promotes intestinal stem cell proliferation through pro-inflammatory cytokine signaling in *Drosophila melanogaster*"

### Supplementary Figure 1

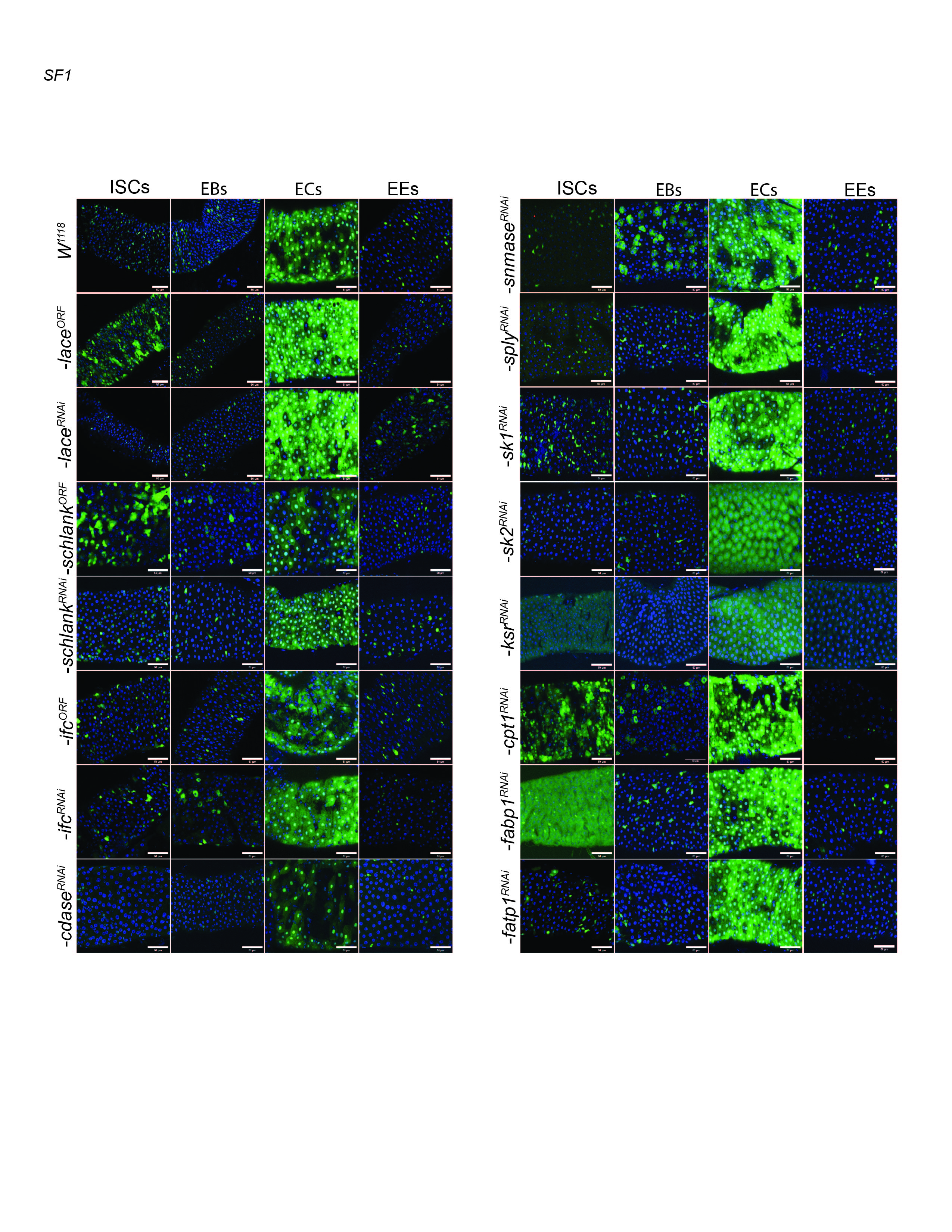

### Supplementary Figure 2

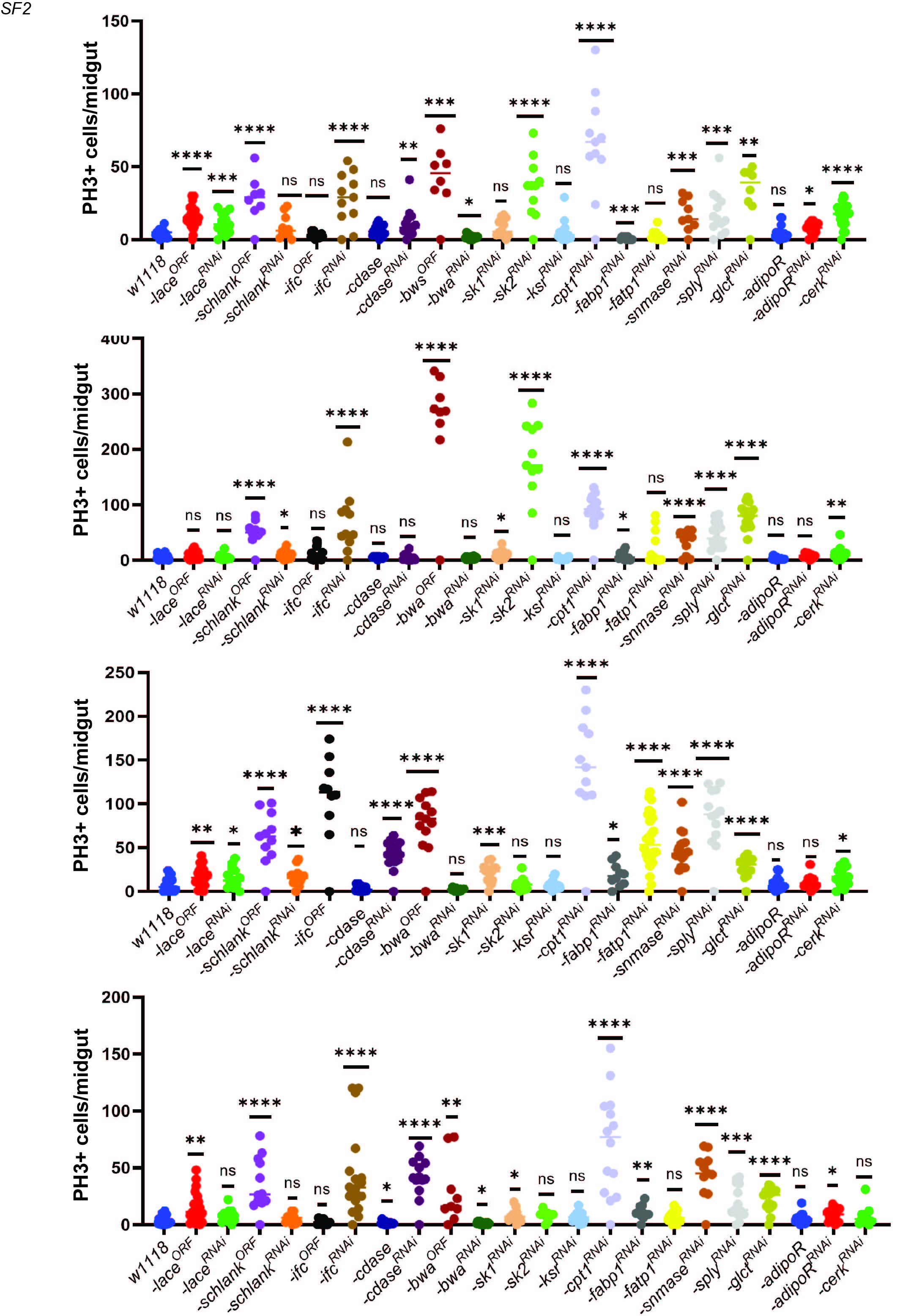

### Supplementary Figure 3

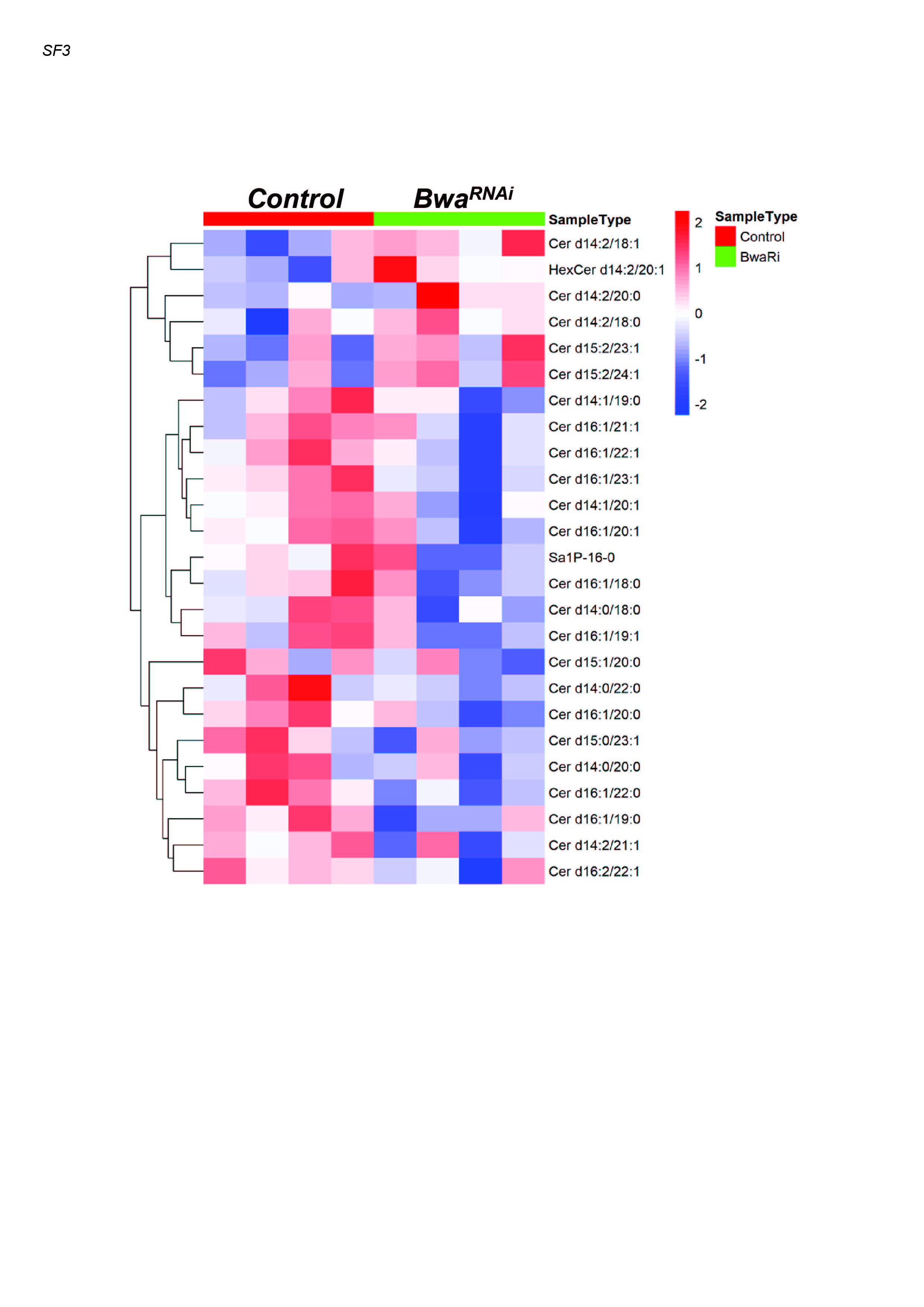

### Supplementary Figure 4

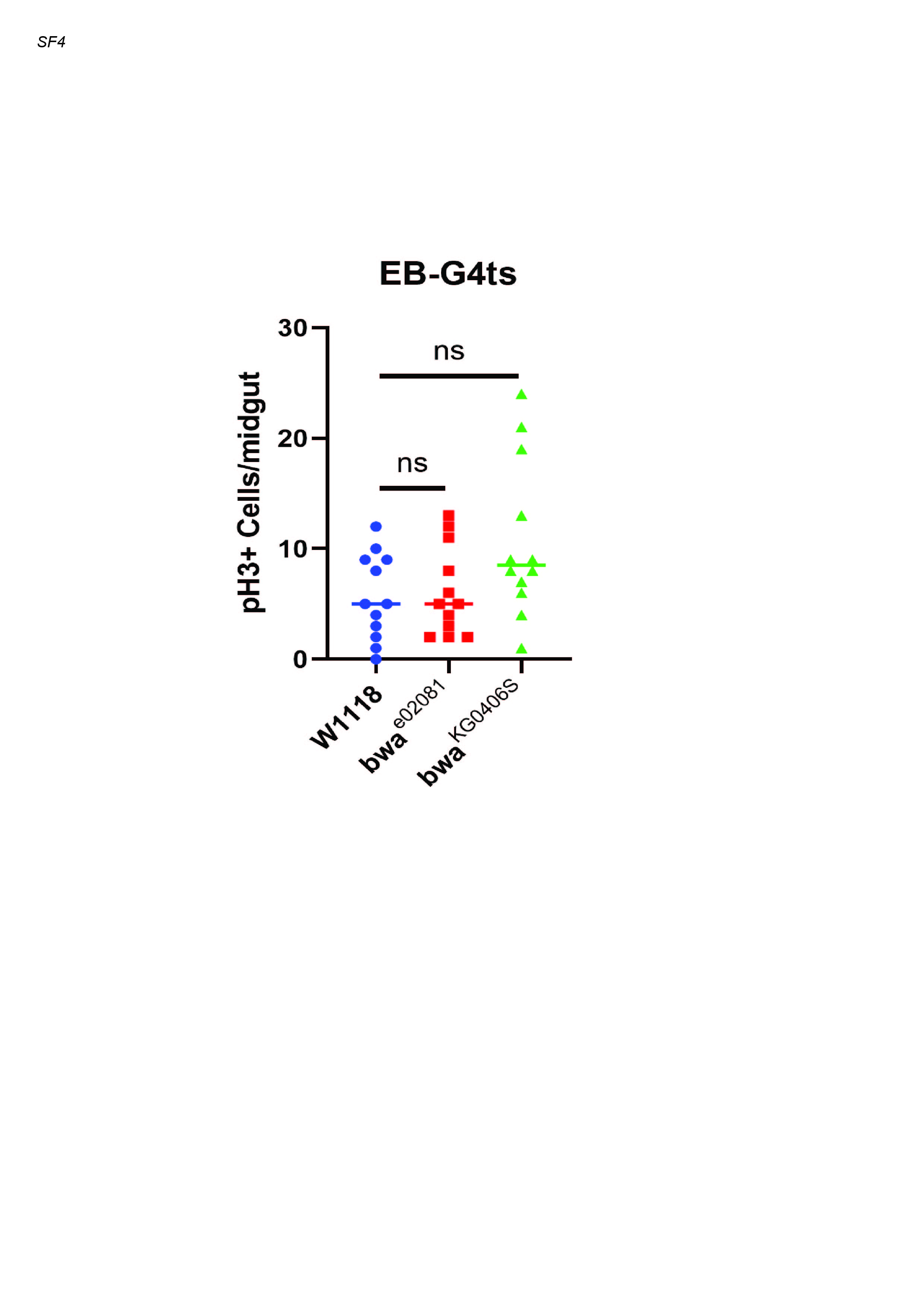

### Supplementary Figure 5

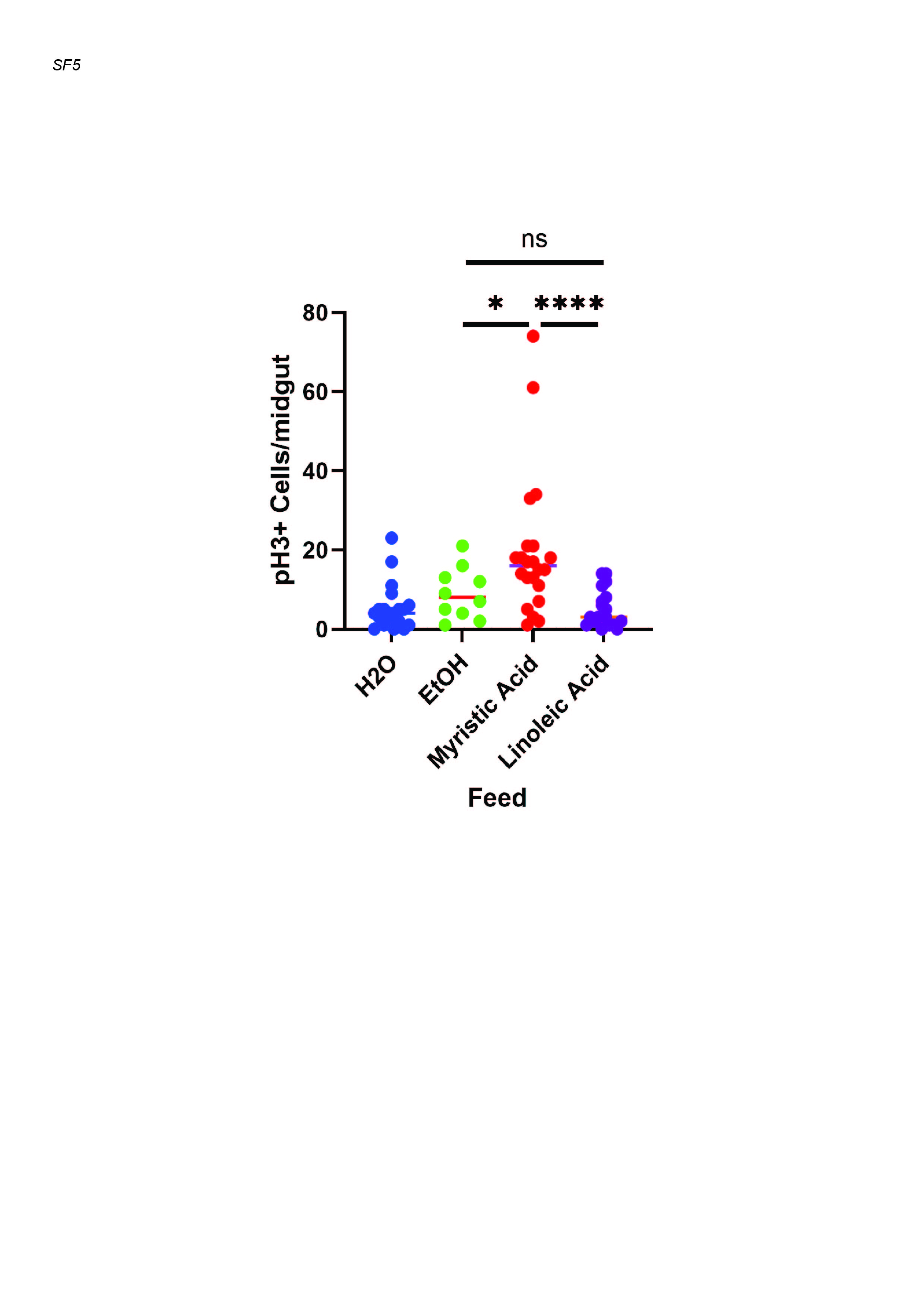

### Supplementary Figure 6

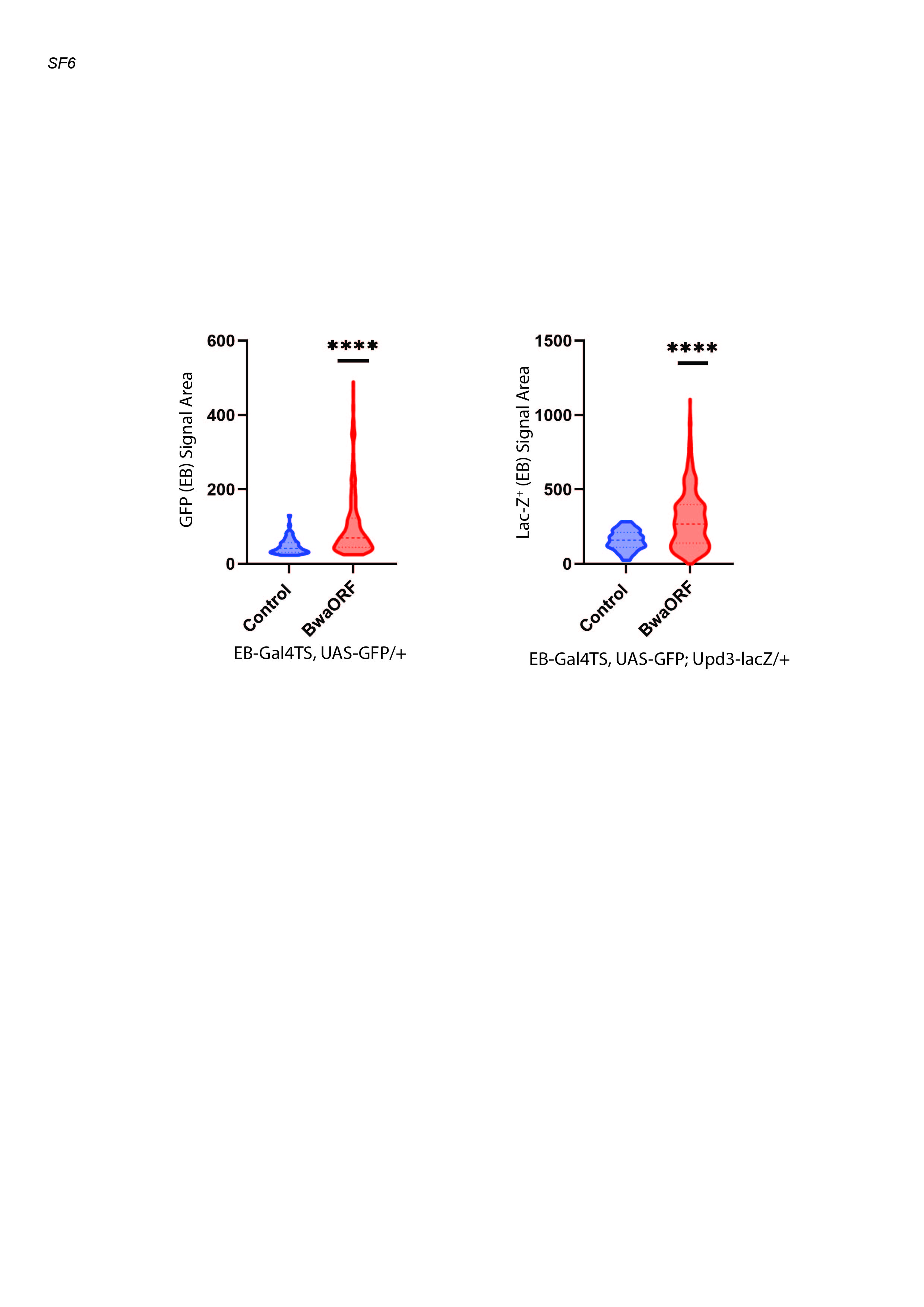
